# A CRISPR-Cas9-based system for the dose-dependent study of DNA double-strand breaks sensing and repair

**DOI:** 10.1101/2021.10.21.465387

**Authors:** Jocelyn Coiffard, Sylvain Kumanski, Olivier Santt, Benjamin Pardo, María Moriel-Carretero

## Abstract

The integrity of DNA is put at risk by different lesions, among which double strand breaks (DSBs) occur at low frequency, yet remain one of the most life-threatening harms. The study of DSB repair requests tools provoking their accumulation, and include the use of chemical genotoxins, ionizing radiations or the expression of sequence-specific nucleases. While genotoxins and irradiation allow for dose-dependent studies, nuclease expression permits assessments at precise locations. In this work, we have exploited the repetitiveness of the Ty transposon elements in the genome of *Saccharomyces cerevisiae* and the cutting activity of the RNA-guided Cas9 nuclease to create a tool that combines sequence specificity and dose-dependency. In particular, we can achieve the controlled induction of 0, 1, 15 or 59 DSBs in cells with an otherwise identical genetic background. We make the first application of this tool to better understand the behavior of the apical kinase of the DNA damage response Tel1 in the nuclear space. We found that Tel1 is capable of forming nuclear foci, which are clustered by condensing when DSBs occur in Ty elements. In striking contrast with other DSB-related protein foci, Tel1 foci are in tight contact with the nuclear periphery, therefore suggesting a role for the nuclear membrane in their congregation.

## Introduction

Cells are confronted to an enormous amount of spontaneous DNA damage arising from natural sources, such as reactive oxygen, reactive carbonyl and nitrogen species, products of lipid peroxidation or the spontaneous chemical lability of the DNA. In addition, there are external sources of damage such as ionizing radiations, chemicals in food, air and water as well as ultraviolet light. Last, rare exposure as that occurring during chemotherapy and radiotherapy is intentionally aimed at forming damage to the DNA. The lesions resulting from these attacks are of a heterogeneous nature, and can include nucleotide base opening, adducts, crosslinks and single-stranded DNA breaks. DNA double strand breaks (DSBs) can also occur, albeit in lower proportion, yet remaining one of the most life-threatening lesions (1).

This higher deleteriousness is in great part due to the fact that, contrary to single strand breaks, DSBs more frequently lack an appropriate undamaged, complementary strand to exploit as a repair template. In fact, DSB repair can efficiently occur without such a support through the process of Non-Homologous End Joining (NHEJ) by sealing the broken ends back together (2). This process could be error-prone because short deletions or insertions may occur at junctions, and translocations take place by the joining between two ends from different DSBs (2). Alternatively, if a sequence similar to the broken one exists elsewhere in the genome, it can be used as a template to guide the information retrieval needed to reconstitute the interrupted sequence at the DSB. This process is Homologous Recombination (HR) and, like NHEJ, is not without drawbacks. The homologous sequence needed to support repair may be available in the replicated sister chromatid or in the homologous chromosome. Further, some sequences are repeated and spread throughout the genome. Given that the resolution of the homology copy may end up with an exchange of the neighboring sequences, a process known as crossing-over, repair involving ectopic repeated sequences or the homologous chromosome could respectively lead to genome rearrangements or loss of heterozygosity and thus only the repair with the sister chromatid warrants a faithful repair (2).

The study of DSB repair necessitates of tools capable of provoking their accumulation. In this sense, genotoxic agents of chemical nature or different types of ionizing radiation are used to damage the genomes of model organisms in a sequence-unspecific manner. A more controlled strategy relies on the expression of proteins whose enzymatic activity creates DSBs at specific DNA target sequences. These can be endonucleases such as restriction enzymes, commonly used *in vitro* for molecular biology (3–8), and meganucleases known to bear a well-defined, rare sequence only present from once to a few times in a given genome (9–13). For example, much of our understanding about DSB repair mechanisms comes from the analysis of HO-mediated DSB in yeast *Saccharomyces cerevisiae* (11). On the one hand, the use of standard restriction enzymes permits the creation of a relatively high number of breaks, at known positions, with ends of a well-defined structure. On the other hand, the use of specific meganucleases cutting at single locations allows to focus the study of molecular events with a high degree of precision while ensuring that no break occurring elsewhere influences the outcome of that DSB event. Yet, a limitation of these sequence-targeted tools is that, contrary to the effects obtained when using increasing doses of genotoxins, they cannot be used for dose-dependent studies. In this sense, in a recent elegant study, Gnügge and Symington engineered yeast strains in which the expression of restriction enzymes could be induced, using a battery of enzymes that can cut an increasing number of sites in the genome, ranging from 20 to 96 (14). However, the different *in vivo* enzymatic activities manifested by each enzyme prevent the use of this system for dose-dependent comparative studies. An alternative work exploited the repetitiveness of the transposable Ty elements in *S. cerevisiae* genome to insert 2, 7 or 11 sites that can be cut upon controlled induction of the HO endonuclease (15). Yet, engineering this system was tedious because it implied multiple rounds of cloning, retrotransposition and Southern blot analysis. This system was used to assess molecular events in DSB repair such as resection and checkpoint activation, as monitored by Southern and western blot, respectively (15,16), and the sensitivity of these techniques allowed to assess dose-dependent differences. Yet, if less sensitive techniques important in the field of DSB sensing and repair need to be used, a system with such a restricted number of breaks may not be sufficient.

In this work, we have exploited the repetitiveness of the Ty transposon elements in the genome of *S. cerevisiae* and the guide RNA-driven sequence specificity of Cas9 cutting activity. In particular, we have targeted an increasing number of Ty elements by designing specific guide RNAs (gRNAs) that could recognize one or several Ty classes. Upon the controlled induction of Cas9 expression, we can achieve an increasing number of enzymatic DSBs (0, 1, 15 or 59) *in vivo*. Because Cas9 and the gRNAs are expressed from plasmids, DSBs can be easily induced in cells grown in otherwise identical conditions and with an identical genetic background. Thus, we have generated a versatile tool that overcomes the lack of dose-dependency of expressing restriction enzymes while achieving the maximum number of DSBs in an easily applicable manner, atypical in systems using sequence-specific nucleases.

We applied this tool to assess the behavior of the Tel1 apical kinase of the DNA damage response in space and time after DSB induction. Contrary to later-acting factors of the DSB repair cascade, fluorescence microscopy is not commonly used to study the very early-acting factors, especially those involved in DSB sensing. Here, we report that Tel1 molecules congregate in the shape of foci in response to diverse sources of DSBs. We report that Tel1 can form up to 8 foci per cell, in striking contrast with other DSB-related proteins, and they can be clustered by condensin when DSBs occur in Ty repetitive elements. We also show that Tel1 foci manifest in tight contact with the nuclear periphery, therefore suggesting a role for the nuclear membrane in their congregation.

## Materials and Methods

*Reagents* used in this work for cell treatments were zeocin, R25001 ThermoFisher Scientific; camptothecin (CPT), C9911 Sigma-Aldrich; and DAPI, D9542 Sigma-Aldrich.

## Culture and treatments

*Saccharomyces cerevisiae* cells carrying plasmids for the expression of the gRNA used to cut the genome and for the expression of the Cas9 endonuclease were grown at 25°C in the appropriate selection medium (-uracil, -leucine). Typically, cells were grown in 2 % glucose; prior to the induction of Cas9 expression, cells were shifted to 2 % glycerol and grown overnight at 25°C to ensure complete glucose consumption. When DNA damage was induced using zeocin (100 µg/mL), cells bearing a control plasmid were grown at 25°C in minimal medium with appropriate selection (uracil). Spot assays were carried out in medium selecting the gRNA plasmids (-uracil) and the inducible Cas9 expression vector (-tryptophan) containing either 2 % glucose or 2 % galactose as a carbon source. Sensitivity spot assays were carried out using YEPD medium supplemented with either DMSO (control) or with 40 µM CPT. For Nup57 tagging with the red fluorophore tDIMER at its genomic *locus*, the plasmid pRS305-*NUP57*-tDIMER (17) (gift from O. Gadal, Toulouse, France) was linearized with *Bgl*II and transformed into strain MM-144. Pus1 was tagged with mCherry using the plasmid YIplac211-mCherry-*PUS1* (18) (a gift from S. Siniossoglou, Cambridge, UK), which was linearized at the *URA3 locus* with *Bgl*II and inserted by HR in this same *locus* in the strain of interest. To accomplish experiments with the *smc2-8* mutant, cultures were split 2h prior to the experiment and cells incubated at the permissive (24°C) or the restrictive (37°C) temperature.

## Pulsed Field Gel Electrophoresis (PFGE)

Agarose plugs containing chromosomal DNA were made as described (19). Chromosomes were separated at 13°C in a 0.9% agarose gel in 0.5× TBE using a Rotaphor apparatus (Biometra) with the following parameters: interval from 100 to 10 s (logarithmic), angle from 120 to 110° (linear), and voltage from 200 to 150 V (logarithmic) during 24 h. The gel was subsequently stained with ethidium bromide for 1 h and washed in water for 30 minutes, then photographed under UV light. For subsequent Southern blot, DNA from gels was transferred to Genescreen Plus membranes (Perkin Elmer). Hybridization was achieved using multiple radioactive probes specific for:

- chromosome III:

> *ARS307* locus; PCR primers 5’-ATTCATTGCGTCTCTGTATTT-3’ and 5’-TTTGAAGATCCTATAACCGTG-3’.

- chromosome IV:

> *SLX5* locus; 5’-GGTAATAACCAAGTCGAAATTTGC-3’ and 5’-CAAAGCACAATTGCACCTCTGATC-3’.

> *FOB1* locus; 5’-CATTTAGTCAAACGGGTGTTA-3’ and 5’-AAATTTTGGAGCATTCCTCTG-3’.

> *RAD9* locus; 5’-GCAGCTCCCCATCAAAATAAG-3’ and 5’-TGAATCTCGTTATTGCTCCTT -3’.

> *MUS81* locus; 5’-AACTTTTTCAGTTTTTGTCGTAATG-3’ and 5’-GGGATGACTATATTTCAAATTGCTA-3’.

> - Ty1 and Ty2; PCR primers 5’-AAATCTGCAAGACAACATGC-3’ and 5’-ATCCTATTACATTATCAATCC-3’.

Hybridized membranes were read using a PhosphorImager (Typhoon IP, GE). Three independent biological replicates were performed for Cas9 induction time course experiments.

### Fluorescence Microscopy

1 mL of the culture of interest was centrifuged, the supernatant thrown away and the pellet resuspended in the remaining 50 µL. 3 µL of this cell suspension were directly mounted on a microscope slide for immediate imaging of the pertinent fluorophore-tagged protein signals. Imaging was achieved using a Zeiss Axioimager Z2 microscope and visualization, co-localization and analysis done using Image J. For time-lapse microscopy experiments, we used cells bearing the x59-cuts gRNA and the inducible Cas9 plasmids that had been induced for Cas9 expression in liquid culture by addition of galactose for 5h40. A total of 10 µL were then spotted on the bottom of a previously prepared FluoroDish (selective minimal medium supplemented with 2% galactose and 1% agarose was casted on FluoroDishes [FD35-100; World Precision Instruments] that had been previously treated with a 0.1% [wt/vol] polylysine solution for 10 min at room temperature, then air-dried). Samples were observed in a Metamorph-controlled Nikon TIRF PALM STORM microscope in a culture chamber at 30°C. Lasers were used at 5% of their power, and pictures were shot every 20 min for 140 min.

### Telomere length measurement

Telomere length was measured by PCR after end labeling with terminal transferase (20,21). End-labeling reactions (40 µL) contained 120 ng genomic DNA, x1 New England Biolabs™ Terminal Transferase Buffer, 1 mM dCTP, 4 units Terminal Transferase (New England Biolabs™) and were carried out at 37°C for 30 minutes followed by heat inactivation at 75°C for 10 minutes. 1/5th volume of 5 M NaCl, 1/80th volume of 1 M MgCl₂ and 1 volume of isopropanol were added to the reaction and DNA was precipitated by centrifugation at 17000×g during 15 min. Precipitated DNA was resuspended in 40 µL of ddH₂O. The end-labeled molecules were amplified by PCR using the primer 5’-GCGGATCCGGGGGGGGGGGGGGGGGG-3’ and 5’-TGTGGTGGTGGGATTAGAGTGGTAG-3’ (X) and 5’-TTAGGGCTATGTAGAAGTGCTG-3’ (Y’), respectively. PCR reactions (50 µL) contained between 40 ng and 80 ng of DNA, 1x myTaq buffer, and primers 0.4 µM each. Amplification was carried out with 5 U of MyTaq polymerase (Meridian Biosciences®). The conditions were 95°C, 5 minutes; followed by 35 cycles of 95°C, 1 minute; 56°C (Y reaction) / 60°C (X reaction), 20 seconds; 72°C, 5 minutes. Reaction was ended with 5 minutes at 72°C. Samples were visualized in a 2 % agarose gel containing 1× GelRed (Ozyme®).

### Serial dilution spots assays

Exponentially growing cells of the indicated genotype were serially diluted 10-fold and 3 µL of each dilution spotted onto the indicated plates, incubated at 30°C for 2 or 3 days and imaged.

### Analysis of DNA content by flow cytometry

430 µL of culture samples at 10^7^ cells/mL were fixed with 1 mL of 100% ethanol. Cells were centrifuged for 1 minute at 16000 xg and resuspended in 500 µL 50 mM Na-Citrate buffer containing 5 µL of RNase A (10 mg/mL, Euromedex, RB0474) and incubated for 2 hours at 50°C. 6 µL of Proteinase K (Euromedex, EU0090-C) were added and after 1 hour at 50°C, cell aggregates were dissociated by sonication (one 3 s-pulse at 50% potency in a Vibracell 72405 Sonicator). 20 µL of this cell suspension were incubated with 200 µL of 50 mM Na-Citrate buffer containing 4 µg/mL Propidium Iodide (FisherScientific). Data were acquired and analyzed on a Novocyte Express (Novocyte).

### Quantifications, Graphical Representations and Statistical Analyses

The number of nuclei displaying foci of the analyzed proteins as well as the number of foci present per nucleus were determined visually by the experimenter. Counting was not done in a blinded manner, but three different researchers were implicated in the quantification of experiments to challenge reproducibility. GraphPad Prism was used both to plot the graphs and to statistically analyze the data. The mean value of nuclei displaying foci was calculated for each independent experiment, and the SEM (standard error of the mean) was used to inform on inter-experiment variation. The SEM estimates how far the calculated mean is from the real mean of the sample population, while the SD (standard deviation) informs about the dispersion (variability) of the individual values constituting the population from which the mean was drawn. Given that the goal of our error bars was to describe the uncertainty of the true population mean being represented by the sample mean, we chose to plot the SEM. To plot probability distributions, we converted the actual observations from each category (number of foci) into frequencies by dividing by the total number of observations. To establish the cutting efficiency, band intensities were determined from non-saturated Southern blot images using ImageJ. The signals emanating from cut bands were divided by the addition of signals emanating from both cut and uncut bands, and expressed as a percentage.

### Strains, plasmids and oligos

The strains used in this study are presented in Table 1 and were obtained either by classical methods for integration, transformation and crosses, or by CRISPR-Cas9 technology in the case of Tel1 yEGFP-tagging. The plasmids used in this study are presented in Table 2. The relevant sequences for CRISPR-Cas9 manipulations are described in Table 3. We have also designed, but not characterized, guide RNAs targeting 2 Ty4, 3 Ty3, 44 Ty1 and 163 Delta elements. Plasmids and sequences are available upon request.

**Table 1.** Strains used in this study.

| Simplified Genotype | Full Genotype | Source |
| --- | --- | --- |
| WT (W303) | *MAT a, ade2-1, his3-11,15, can1-100, leu2-3,112, trp1-1, ura3-1, GAL+, psi+, RAD5* | PP870, Philippe Pasero |
| *tel1Δ* | *MAT a, ura3-52, leu2-3,112, his3-i200, trp1-1, trp1-1, lys2-801, tel1ΔKAN<sup>R</sup>* | PP1217, Philippe Pasero |
| Rad52-YFP<br>Rfa1-CFP<br>mCherry-Pus1 | *MAT a, ADE2, his3-11,15, can1-100, leu2-3,112, trp1-1, ura3-1, RAD52-YFP*<br>*RFA1-CFP ura3::mCherry-PUS1::URA3* | PP3558, Philippe Pasero |
| yEGFP-Tel1<br>mCherry-Pus1 | *MAT a, ade2-1, his3-11,15, can1-100, leu2-3,112, trp1-1, ura3-1, GAL+, psi+, RAD5, yEGFP-TEL1, ura3::mCherry-PUS1::URA3* | MM-40, this study |
| yEGFP-Tel1 | *MAT a, ade2-1, his3-11,15, can1-100, leu2-3,112, trp1-1, ura3-1, GAL+, psi+, RAD5, yEGFP-TEL1* | MM-144, this study |
| yEGFP-Tel1<br>Nup57-tDimer | *MAT a, ade2-1, his3-11,15, can1-100, leu2-3,112, trp1-1, ura3-1, GAL+, psi+, RAD5, yEGFP-TEL1, NUP57-tDIMER-RFP::LEU2* | MM-282, this study |
| *smc2-8* yEGFP-Tel1 | *MAT a, ade2-1, his3-11,15, can1-100, leu2-3,112, trp1-1, ura3-1, GAL+, psi+, RAD5, yEGFP-TEL1, smc2-8::KAN<sup>R</sup>* | MM-434, this study |
| *lig4Δ* | *MAT a, ade2-1, his3-11,15, can1-100, leu2-3,112, trp1-1, ura3-1, GAL+, psi+, RAD5, lig4Δ::KAN<sup>R</sup>* | PP3213, Philippe Pasero |
| *rad52Δ* | *MAT a, ade2-1, his3-11,15, can1-100, leu2-3,112, trp1-1, ura3-1, GAL+, psi+, RAD5, rad52Δ::klTRP1* | PP6333, Philippe Pasero |

**Table 2.**
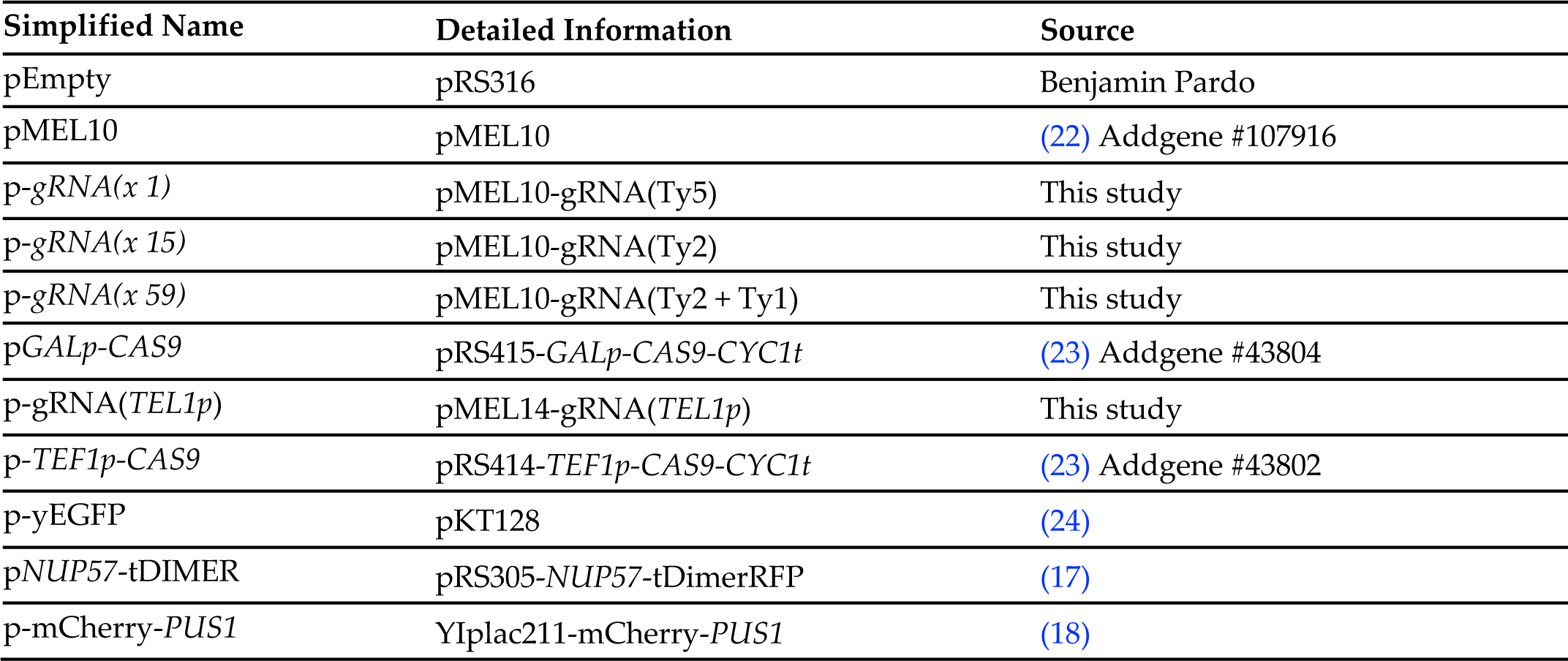
Plasmids used in this study.

| Simplified Name | Detailed Information | Source |
| --- | --- | --- |
| pEmpty | pRS316 | Benjamin Pardo |
| pMEL10 | pMEL10 | (22) Addgene #107916 |
| p-gRNA(*x 1*) | pMEL10-gRNA(Ty5) | This study |
| p-gRNA(*x 15*) | pMEL10-gRNA(Ty2) | This study |
| p-gRNA(*x 59*) | pMEL10-gRNA(Ty2 + Ty1) | This study |
| pGALp-CAS9 | pRS415-GALp-CAS9-CYC1t | (23) Addgene #43804 |
| p-gRNA(TEL1p) | pMEL14-gRNA(TEL1p) | This study |
| p-TEF1p-CAS9 | pRS414-TEF1p-CAS9-CYC1t | (23) Addgene #43802 |
| p-yEGFP | pKT128 | (24) |
| pNUP57-tDIMER | pRS305-NUP57-tDimerRFP | (17) |
| p-mCherry-PUS1 | YIplac211-mCherry-PUS1 | (18) |

**Table 3.**
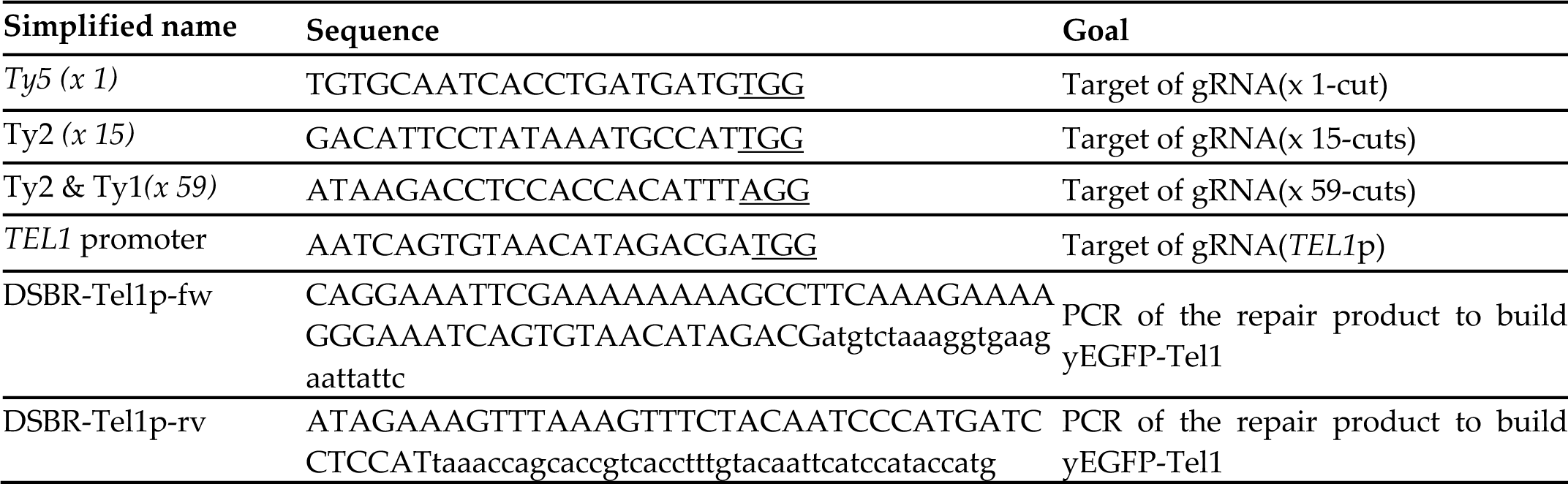
Relevant sequences used in this study (PAM is underlined)

## Results

### A genetic system to create DNA double strand breaks in an inducible and dose-dependent manner

In order to generate multiple DSBs, we targeted repeated DNA by using the CRISPR-Cas9 technology (25). Among the most abundant classes of repeats in *S. cerevisiae* are retrotransposons (Ty elements), which represent about 3% of the genome (26), and whose targetability by the CRISPR-Cas9 technology has already been demonstrated in a study aimed at engineering translocations in a controlled manner (27). Based on the sequenced genome of the W303 yeast strain (28), we designed guide RNAs (gRNAs) to target the ORF of the sole complete Ty5 element (1 targeted site), of the 15 copies of Ty2 (15 targeted sites) and of both the closely related Ty1 and Ty2 (59 targeted sites) (Fig 1A). The sequences for the gRNAs were cloned in multicopy expression plasmids, different from the one expressing the Cas9 nuclease under the control of the galactose-inducible GalL promoter (22,23).

**Figure 1.**
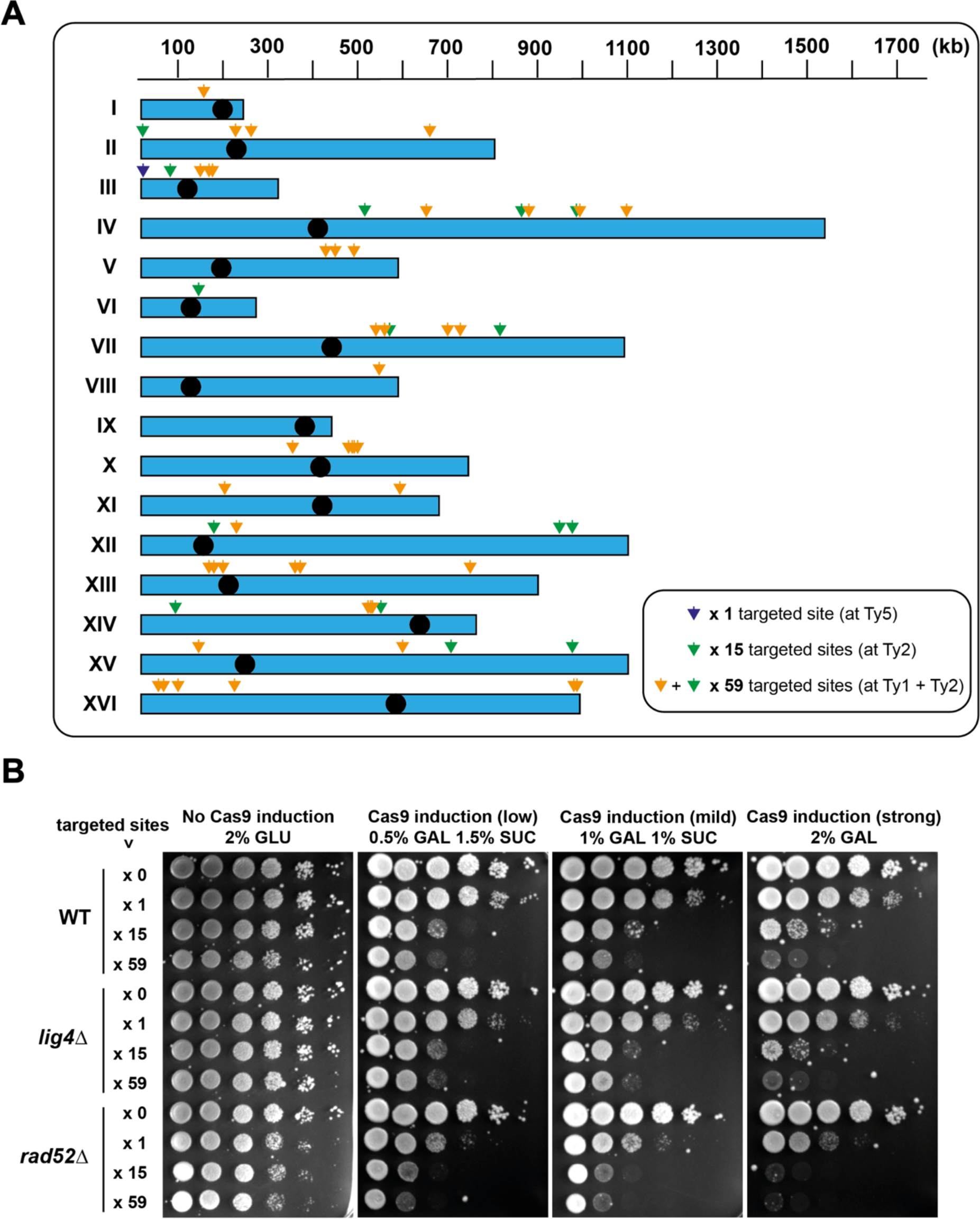
Design of the CRISPR-Cas9-based system to induce DSBs in a dose-dependent manner. **A.** Scheme of the sixteen *S. cerevisiae* chromosomes (blue boxes), in which the black circles mark the relative position of the centromeres. Color-coded arrows mark the approximate Ty positions targeted by the designed gRNAs. In more detail, the x 1 gRNA targets the single Ty5 (dark blue arrow), the x 15 gRNA targets the Ty2 sites (green arrows), and the x 59 gRNA targets both Ty2 and Ty1 sites (orange and green arrows). **B.** *S. cerevisiae* wild-type, *lig4Δ* and *rad52Δ* cells were transformed with the vector bearing an inducible Cas9 and with the vector expressing the relevant gRNA to achieve the desired number of breaks. 10-fold serial dilutions of cells exponentially growing in glucose to prevent Cas9 expression were spotted onto selective minimal medium plates supplemented with the following carbon source: either glucose, to monitor the loading control; or increasing amounts of galactose complemented with sucrose to modulate Cas9 expression. Growth plates were incubated 3 days at 30°C and imaged. The experiment was repeated three times.

To assess the performance of the system, we initially tested viability upon inducing Cas9 expression in wild-type cells simultaneously expressing no guide RNA (gRNA) and thus not inducing DSBs (x 0), or the gRNAs leading to the potential cleavage of the genome at 1 site (x 1), 15 sites (x 15) or 59 sites (x 59), respectively. We spotted serial dilutions of cells either in medium containing glucose (no Cas9 expression) or increasing concentrations of galactose (low, mild and strong Cas9 expression). We observed that the viability of wild-type cells expressing x 1 gRNA was only mildly affected compared with no guide RNA (x 0). This result is not surprising taking into account that the x 1 gRNA is targeting Cas9 to the sole Ty5 element, which is located very close to the left telomere of chromosome III. Cleavage of chromosome III at this location generates a telomere-proximal fragment that does not contain essential genes and could be lost, while the other fragment containing the centromere could be efficiently repaired by *de novo* telomere addition or telomere recombination. In contrast to the effect of the x 1 gRNA, the viability of wild-type cells expressing x 15 or x 59 gRNAs was strongly affected by the increased number of targeted sites (x 0 < x 1 < x 15 < x 59) and by the increased expression of Cas9 (Fig 1B, WT). These results suggest that DSBs are readily induced by the expression of Cas9 in a dose-dependent manner.

Our next objective was to provide a physical and more quantitative characterization of the system. To this end, we grew wild-type cells overnight until they reached exponential phase in medium selective for the Cas9 and gRNAs plasmids and with glycerol as the carbon source, allowing a robust and controlled induction of Cas9 upon galactose addition. Cells were recovered and processed for flow cytometry and Pulsed Field Gel Electrophoresis (PFGE) analyses before induction of Cas9 expression and every 2 hours thereafter during 8 hours. The analysis of DNA contents by flow cytometry showed a mild but progressive accumulation of cells in G_2_/M in the x 15 and x 59 systems (Fig 2A, red asterisks). While these results are consistent with the activation of the DNA damage checkpoint, which halts the cell cycle progression in G_2_/M in response to DSBs (29), the modest cell cycle halt suggests either low cleavage efficiency and / or fast repair (see below).

**Fig 2.**
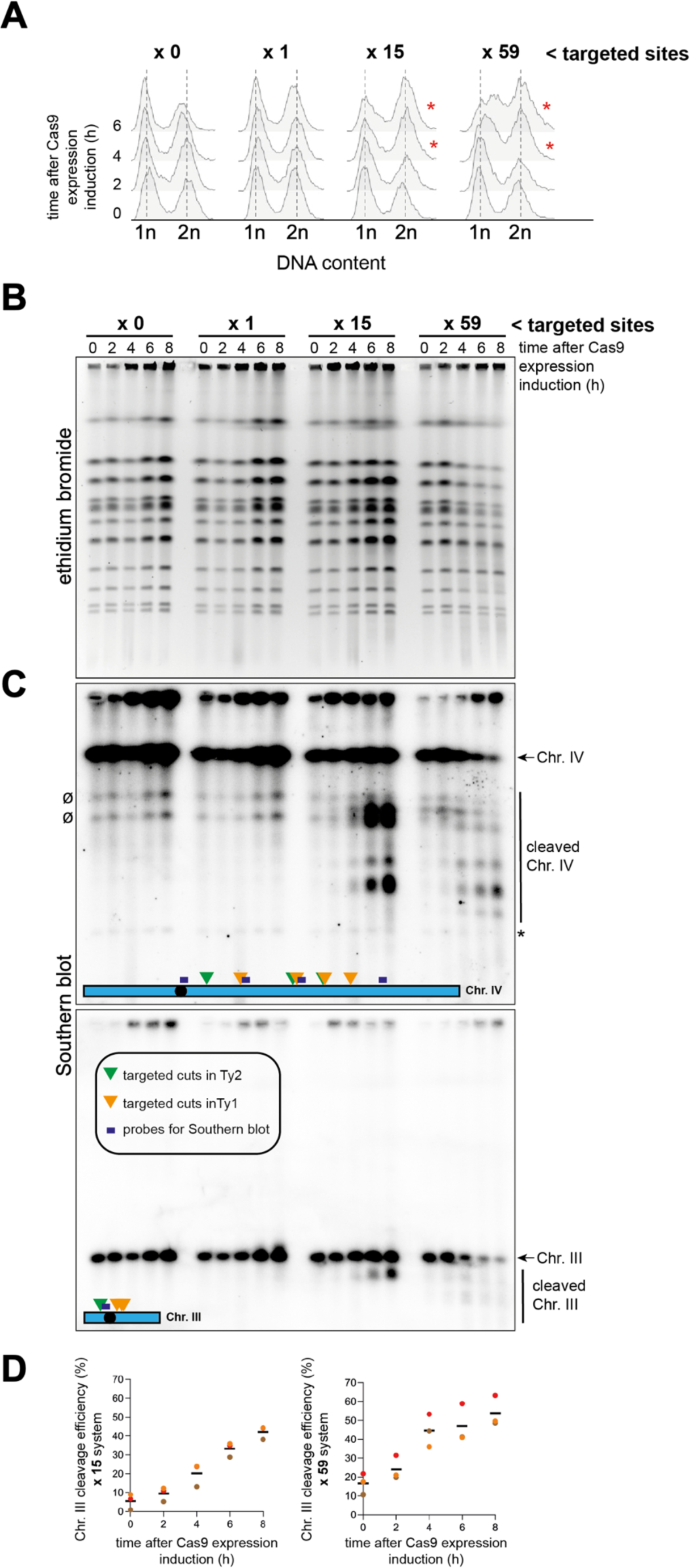
Physical characterization of the DSB-inducible system. **A.** *S. cerevisiae* wild-type cells were transformed as in Fig 1B and a culture of cells growing exponentially in selective minimal medium with glycerol as the carbon source was prepared. A sample was taken at time 0 (before induction). After the addition of 2% galactose, samples were taken at the indicated time points to assess cytometry profiles. n and 2n refer to the DNA content, thus serving as an estimate of the number of cells in G_1_ and G_2_ phases of the cell cycle, respectively. The red asterisks indicate the time-points at which an increase in the number of cells in G_2_/M is detected when compared to the no-cut condition. **B.** *S. cerevisiae* wild-type cells were transformed as in Fig 1B, and a culture of cells growing exponentially in selective minimal medium with glycerol as the carbon source was prepared. A sample was taken at time 0 (before induction). After the addition of 2% galactose, samples were taken at the indicated time points. Cells were processed for Pulsed Field Gel Electrophoresis (PFGE). This technique allows for the separation of chromosomes. The PFGE gel was dyed with ethidium bromide. **C.** Southern blot against the DNA from the PFGE shown in **(B)** using one probe targeting chromosome III and four probes targeting chromosome IV. Bands corresponding to full-length chromosomes III and IV are indicated. The asterisk indicates a remaining band corresponding to chromosome III after incomplete stripping of the radioactive probe from the membrane, while the symbol Ø points at unspecific signals present at all time-points. **D.** Quantification of chromosome III cleavage efficiency after Cas9 induction in the x 15 and the x 59 systems. The mean values (dark bar) and the individual values (circles) from three independent experiments (indicated by different colours) are plotted for each time point of the time course experiment shown in **(C)**.

Separation of chromosomes by PFGE allows for the detection of broken DNA molecules. No broken DNA molecules could be detected at any of the time-points of the x 0 kinetics. However, modest smeared signals were visible in the ethidium bromide-stained gels at late times of the x 15 and x 59 gRNA kinetics (Fig 2B). Yet, the strongest evidence of molecules being broken emanated from the latest time-points in the x 59 system, when signals corresponding to full chromosomes started to fade away. To provide formal proof of the breaks, the DNA was transferred and the membranes subjected to Southern blotting using either a probe directed to the chromosome III or a mix of four probes directed against several locations distributed between the potential break sites on the chromosome IV. As a result, a time- and dose-dependent pattern of broken chromosome fragments could be observed (Fig 2C), confirming the proficiency of the system. Analysis of the fragments released from chromosome III in the x 15 and x 59 systems allowed us to quantify Cas9’s cleavage efficiency. It revealed a progressive increase over time reaching mean values between 43 and 54% 8 h after Cas9 induction (Fig 2D). We conclude that the CRISPR-Cas9-based system produces DSBs with a good efficiency.

We subsequently conducted a deep restriction analysis of chromosome cleavage in order to verify whether Cas9 could cleave all the potential cut sites in Ty elements, as indicated in Fig 1A. We used the known sizes of the full-length chromosome bands from the ethidium bromide-stained gel to infer the sizes of the chromosome fragments that appeared 8h after Cas9 induction on the Southern blot membrane (S1 Fig). We could not perform this analysis with the x 1-cut system since cleavage of chromosome III should happen very close to the telomere and generate a fragment which size is not distinguishable from the full-length chromosome after separation by PFGE. In the x 15 system, the chromosome III is expected to be cleaved once, generating a ∼254 kb fragment, which we could detect by using a probe targeting this fragment (Fig 3A). In this same system, 3 cuts are expected in chromosome IV. These cuts can generate various fragments corresponding to the complete or partial cleavage of the chromosome IV (Fig 3A). The Southern blot analysis revealed the presence of bands corresponding to all the expected sizes of the chromosome fragments (Fig 3A). We performed the same analysis for the x 59 system, in which four cuts in chromosome III and seven cuts in chromosome IV are expected. Again, we could identify chromosome fragment bands corresponding to all expected restriction patterns thanks to various probes targeting the regions between the cleavage sites (Fig 3B). This analysis thus indicates that Cas9 could efficiently cleave chromosomes III and IV in all targeted regions, inferring that the CRISPR-Cas9-based system is able to efficiently induce DSBs in every chromosome in a site-specific and dose-dependent manner.

**Fig 3.**
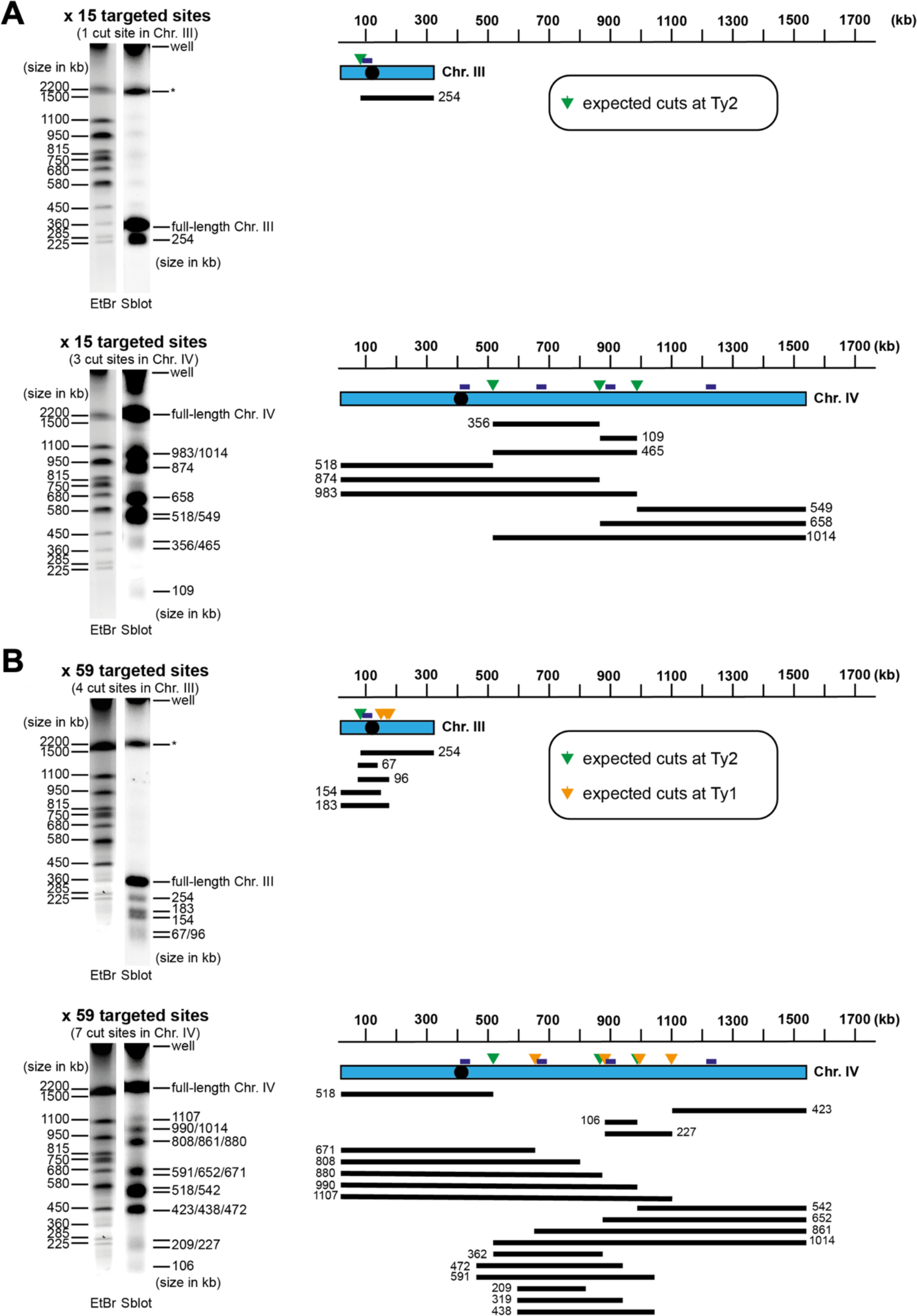
Restriction analyses of chromosome cleavage. **A, B.** Restriction analysis of chromosomes III and IV cleaved by Cas9 in the x 15 **(A)** and x 59 **(B)** systems. Full-length chromosome sizes (from ethidium bromide-stained gel; EtBr) and inferred fragment sizes (from Southern Blot; Sblot) are indicated in kb. The asterisk indicates a remaining band corresponding to chromosome IV after incomplete stripping of the radioactive probe from the membrane. The approximate Ty positions targeted by the designed gRNAs and the probes used for Southern blot analysis are indicated on chromosome schemes. Restriction patterns are depicted, with expected chromosome fragments sizes, out of both full and partial digestions, indicated in kb. Shown images come from the experiment shown in S1 Fig.

### The kinetics of DNA repair foci formation reflects the dose-dependent induction of DSBs by Cas9

DSB repair primarily occurs by Homologous Recombination (HR) or Non-Homologous End Joining (NHEJ). In cycling *S. cerevisiae* cells, repair by HR is generally favored over NHEJ by the availability of the sister-chromatid during DNA replication (2,30). We wondered if the repair of DSBs in repeated Ty elements would follow this general rule. We repeated the viability assay (Fig. 1B) by spotting cells either lacking Lig4, the DNA ligase involved in NHEJ, or Rad52, the master regulator of HR. The loss of viability of *lig4Δ* cells upon Cas9 expression was similar to wild-type cells, indicating that NHEJ is not involved in cell survival to Cas9-induced DSBs. However, the lack of Rad52 in *rad52Δ* cells negatively impacted the viability by about one order of magnitude compared to wild-type cells, even for the 1 x system (Fig. 1B). These results indicate that HR is involved in the repair of Cas9-induced DSBs in Ty elements, as expected for DSB repair outside repeated DNA sequences in cycling cells.

In order to monitor the dynamics of repair by HR, we exported the system (Cas9 and gRNAs plasmids) to an otherwise wild-type strain in which early HR actors are fluorescently tagged. The Rfa1 component of the heterotrimeric RPA complex, binding exposed single stranded DNA (ssDNA) at broken ends, was tagged with CFP at its C-terminus; Rad52, which replaces RPA by Rad51 on ssDNA to initiate homology search, was tagged with YFP at its C-terminus. As above, cells were grown overnight to mid-log phase in selective medium containing glycerol, then galactose was added to induce Cas9 expression. Cells were visualized through a fluorescence microscope and images acquired before Cas9 expression induction and every hour during 7 hours thereafter. We counted the percentage of nuclei in the population displaying Rfa1-CFP and Rad52-YFP foci (at least one focus) along time. The percentage of nuclei basally displaying Rfa1 foci was of 10 % and fluctuated around this value for each time point when Cas9 was expressed in the absence of any gRNA (Fig 4A,B, x 0). Likewise, the lack of DSB induction maintained a basal level of 4 % of the nuclei displaying Rad52-YFP foci (Fig 4A,C, x 0). These data agree with previously reported basal levels of foci for these factors (31), and further indicates that, in the absence of any gRNA, Cas9 alone does not trigger any accumulation of DNA damage. Importantly, the expression of gRNAs driving an increasing number of DSBs permitted us to draw the following observations: First, the initial percentage of foci-displaying cells increased with time when the x 1-, x 15- and x 59 targeted sites gRNAs were present in the cells, confirming the proficiency of the system (Fig 4B,C). Second, the proportion of nuclei displaying Rfa1 foci was consistently double than that of nuclei bearing Rad52 foci, probably reflecting the increased residence time of resected filaments in comparison with the process of homology search (Fig 4B,C). Third, and as a general rule, gRNAs targeting a higher number of sites led to an increased number of cells bearing Rfa1 and Rad52 foci (*i*.*e*. x15 > x1 > x0), with the higher mean values being of 45% of cells bearing Rfa1 foci and 27% showing Rad52 foci (Fig 4B,C). Surprisingly, from 5 hours onwards, creating 15 DSBs in the genome triggered more Rfa1 foci accumulation than inducing 59 DSBs, and a much faster (although equal in value) accumulation of Rad52 foci (Fig 4B,C). This may highlight the phenomenon of interference at clustered DSBs. Indeed, DSBs concentrated at near-by locations, as those induced in the x 59 system, are less efficiently repaired than isolated lesions (32). Finally, we observed that increasing numbers of DSBs also increased the number of individual Rfa1 foci per nucleus (Fig 4D). This was not the case for Rad52 foci, which were previously described to be repair centers capable of recruiting more than one DSB (33). Overall, these results suggest that DSBs induced by Cas9 engage into HR in a dose-dependent manner.

**Fig 4.**
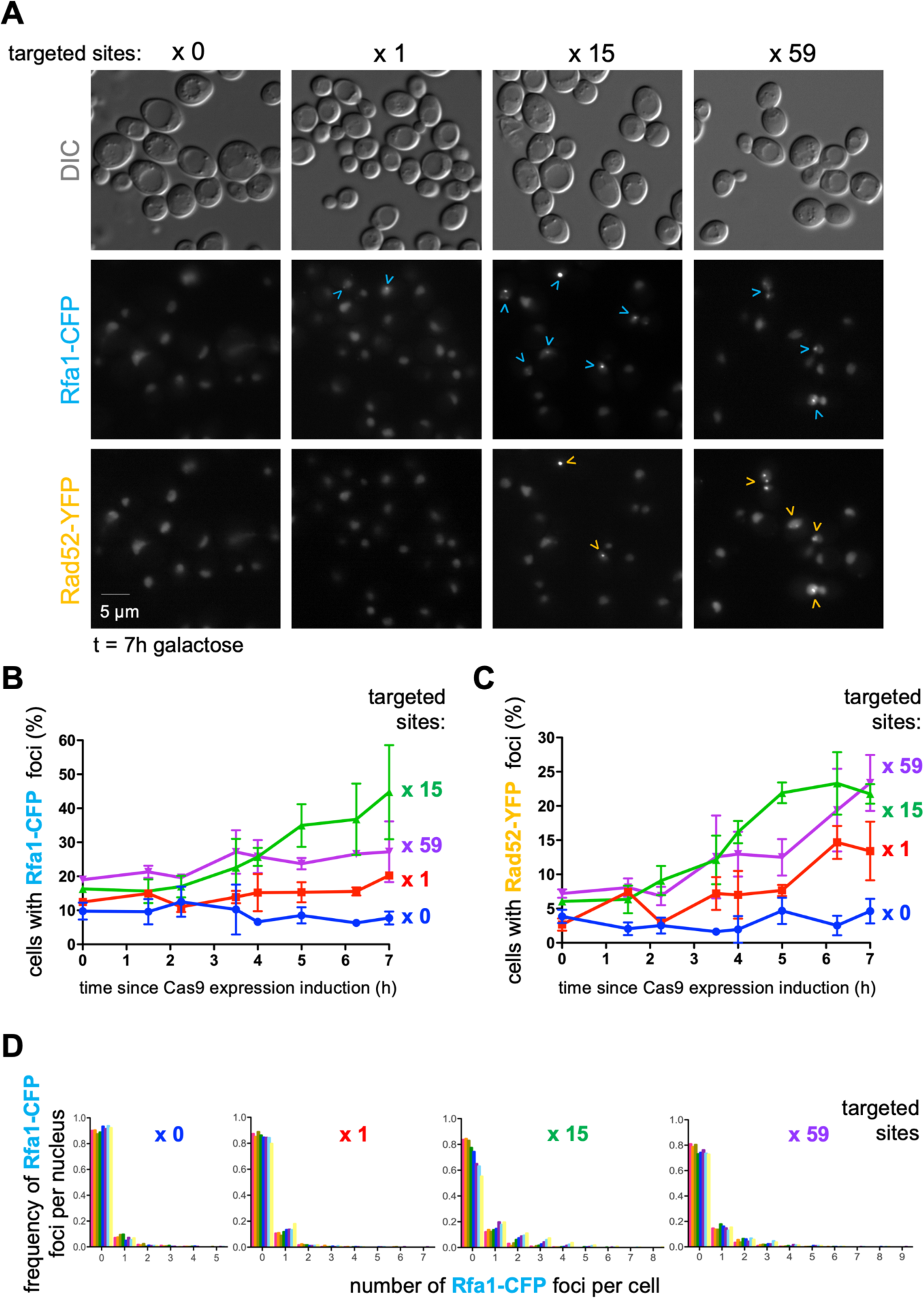
Characterization of DNA repair foci formation in response to increasing Cas9-induced DSBs. **A.** Wild-type cells transformed with the vector expressing an inducible Cas9 and the plasmid expressing the gRNA driving the desired number of cuts grown to exponential phase in selective minimal medium using glycerol as the carbon source. A sample was taken at time 0 before galactose was added to induce Cas9 expression, samples retrieved at the indicated times, and cells inspected by microscopy. Representative images of Differential Interference Contrast (DIC), Rfa1-CFP and Rad52-YFP channels are shown. Arrowheads point at foci formed by the fluorescently tagged proteins. **B.** Percentage of cells in the population displaying at least one focus of Rfa1-CFP in samples from **(A)** at different times since galactose addition. Each point is the mean of three independent experiments, and the error bars represent the SEM out of those three experiments. **C.** Graph showing the percentage of cells in the population displaying at least one focus of Rad52-YFP in samples from **(A)** at different times after galactose addition. Details as in **(B)**. **D.** Graphs showing the frequency of Rfa1-CFP foci in a given nucleus at a given time as calculated from the experiments presented in **(C)**. The probability distribution is calculated upon merging the three experiments presented in **(C)**. All three independent experiments had a similar profile. At least 200 cells were counted per time-point, condition and experiment.

### Time-lapse fluorescence microscopy reveals multiple rounds of Rad52 foci formation in G_2_/M cells

Because some DSBs may be repaired while others are formed, we posited that the frequency of cleavage and the percentage of repair events could be higher than we calculated. To test this hypothesis, we used time-lapse live fluorescence microscopy to look at continuous Rad52 foci formation. Cells bearing the x 59 system were induced for Cas9 expression in liquid medium containing galactose for 5h40, then mounted on an imaging device under an agarose pad prepared in selective minimal medium with galactose to keep Cas9 expression on. 300 and 540 individual cells were monitored in two independent experiments every 20 min for 140 min (therefore reaching 8h post-galactose addition). Illustrative examples of individual cells are shown evolve in time in Fig 5. We observed two types of Rad52-YFP populations contributing differently to Rad52 foci counts. One population of cells displayed only one event of Rad52 foci formation. These events were long-lasting with a duration of at least 60 min and even beyond 140 min (Fig 5). These events readily contribute to the observed percentage of positive cells when performing classical experiments in which multiple pictures of a culture are taken at a given time (as in Fig 4C). Yet, a second population exists in which two events of Rad52 foci formation were observed during the time-lapse experiment. These events were short-lasting, for their duration ranged from less than 20 min to maximum 60 min. In these cells, a lapse of time frequently exists during which no foci can be seen, then new foci appear, indicative of a new round of cut(s) and repair (Fig 5). When displaying Rad52 foci-containing cells according to their cell cycle stage, as estimated from their shape and bud growth, the population bearing long-lasting Rad52 foci were likely early-S phase cells, showing no or small buds. On the contrary, the cells displaying short-lived Rad52 foci corresponded to G_2_/M cells. These results suggest that DSBs induced early during the cell cycle could not be quickly repaired by HR and that the completion of DNA replication may be required for HR to proceed further. Once in G_2_/M, DSBs could undergo fast repair due to the availability of sister chromatids, allowing new rounds of cleavage by Cas9 and Rad52 foci formation. Importantly, since the foci displayed by the same cell could stem from different cutting events, we reckon there is an underestimation of the real cleavage frequency when calculated from snapshots taken every hour, as those shown in Fig 4C. Finally, we analyzed the karyotypes of 46 independent survivor colonies coming from wild-type cells that had endured 8 hours of Cas9 induction with the x 59 system and did not observe any gross chromosomal rearrangements (S2 Fig). We conclude that the repair of the DSBs induced by Cas9 is faithful, despite the fact that DSBs occurs in repeated Ty elements dispersed throughout the genome. These results are consistent with the relatively low frequency of genomic alterations observed after targeting Cas9 to Ty1 elements in haploid yeasts (34).

**Fig 5.**
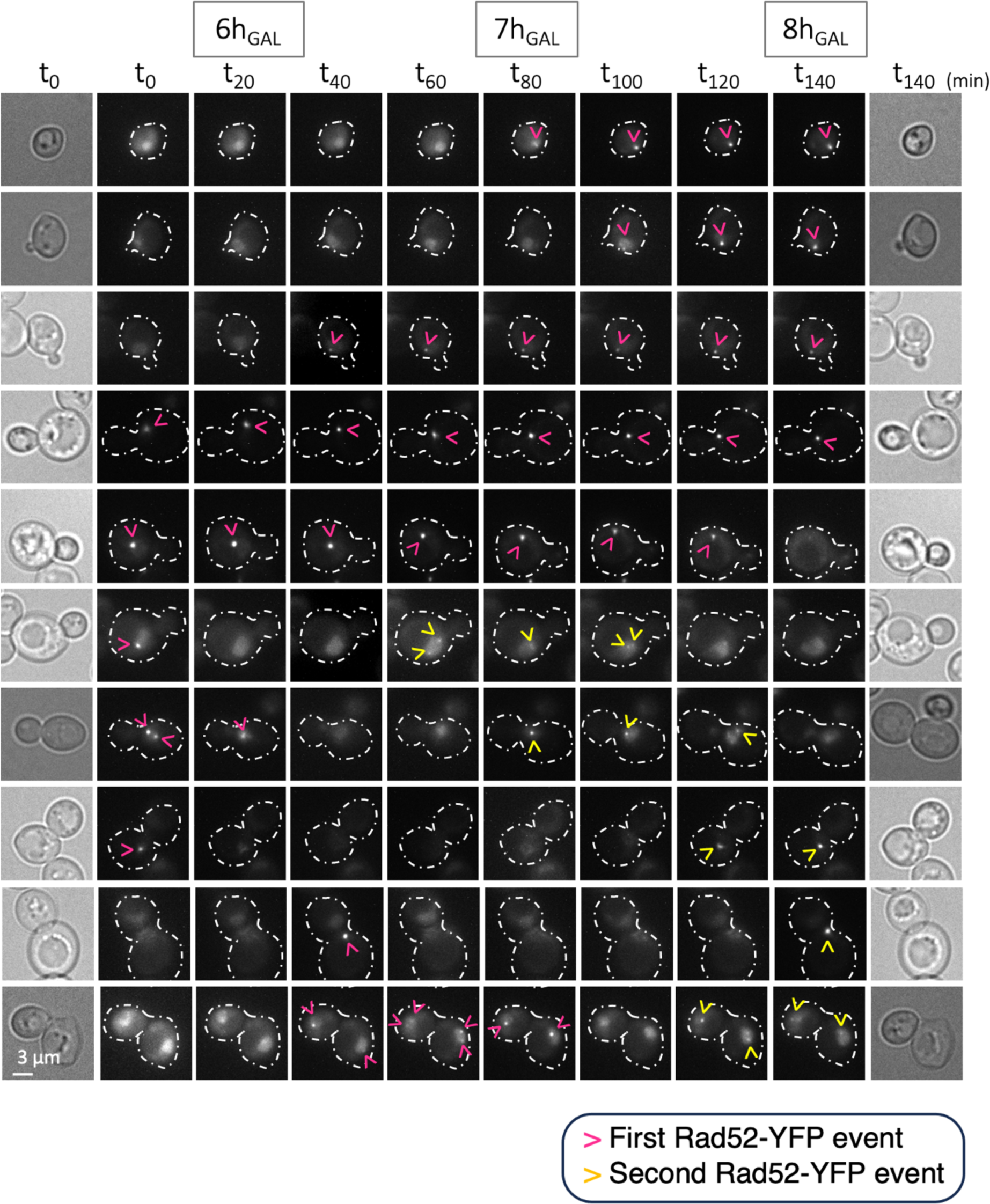
Time-lapse microscopy analysis of Rad52 foci formation after Cas9-induced DSBs. *S. cerevisiae* wild-type cells in which Rad52 was YFP-tagged were transformed with the vector bearing an inducible Cas9 and with the vector expressing the x 59 gRNA, kept in selective medium plus glycerol as the source of carbon, then changed to galactose to induce Cas9 expression. After 5h40, cells were mounted on a microscope device and immobilized using an agarose pad embedded in the same medium. Images of multiple positions were acquired every 20 min for 140 min. Individual cells followed this way are shown evolving in time. The first Rad52-YFP foci to be detected are indicated by pink arrowheads, while secondary events are indicated by yellow ones. The first and last time-point brightfield images are also shown. On the vertical axis, the cells have been intentionally ordered by their morphology, with those in early S-phase at the top and those in late S-phase or G_2_/M at the bottom.

### Exploiting the dose-dependent DSB system to study DSB signaling by Tel1

Fluorophore-tagging of multiple proteins has allowed the establishment of the temporal kinetics from DSB sensing till late steps of its repair (31,35). Yet, compared to the number of works assessing the formation, persistence, dissolution or frequency of foci of DNA repair proteins such as Rad52, the study of very early acting factors such as DSB sensors has been under-assessed. DSBs sensing is orchestrated by the early arrival of the MRX (MRN in humans) complex and its immediate binding by the apical kinase of the DNA Damage Response Tel1 (ATM in humans). Mre11 and Tel1 foci can form at any stage of the cell cycle, persist if resection is not implemented, as in *SAE2* deletion mutants (31,36) and, in agreement with their early role, do not depend on the ssDNA-coating complex RPA (31). Still, in contrast to Mre11 foci, which have attracted more interest (16,36), Tel1 study by microscopy has been assessed in only one study (31). This little interest may relate to the fact that the absence of Tel1 hardly sensitizes cells to genotoxic agents, with the exception of Topoisomerase 1 trapping by camptothecin (CPT) (37), presumably because its deficiency is often compensated by the other apical kinase Mec1 (38,39). Yet, accurate sensing, proper processing and timely checkpoint activation upon DSBs depend on Tel1, and therefore the alternative orchestration by Mec1 may not reflect the physiological pathway that the lesions should have triggered. Moreover, it has been observed that Tel1 signaling becomes critical in the absence of Mec1 in response to an increasing number of simultaneous DSBs (16).

We have tagged Tel1 with yeast-enhanced-GFP (yEGFP) at its N-terminus, a location reported to preserve its function (31,40), while conserving its natural promoter. In agreement, strains bearing this modification were as proficient as their isogenic wild-type in tolerating CPT, in contrast to *tel1Δ* cells (Fig 6A). Tel1 function in DSB signaling is key in preserving telomere length, which can be measured by subjecting genomic DNA samples to terminal transferase treatment followed by PCR-driven telomere amplification (20,21). We monitored both X and Y’ telomere length and observed similar sizes in wild-type and in derived yEGFP-Tel1 cells, in marked contrast with the shorter products observed in *tel1Δ* cells (Fig 6B). Thus, we concluded that the fluorescent yEGFP tag at the N-terminus of Tel1 does not alter its biology and can be used to assess functional questions.

**Fig 6.**
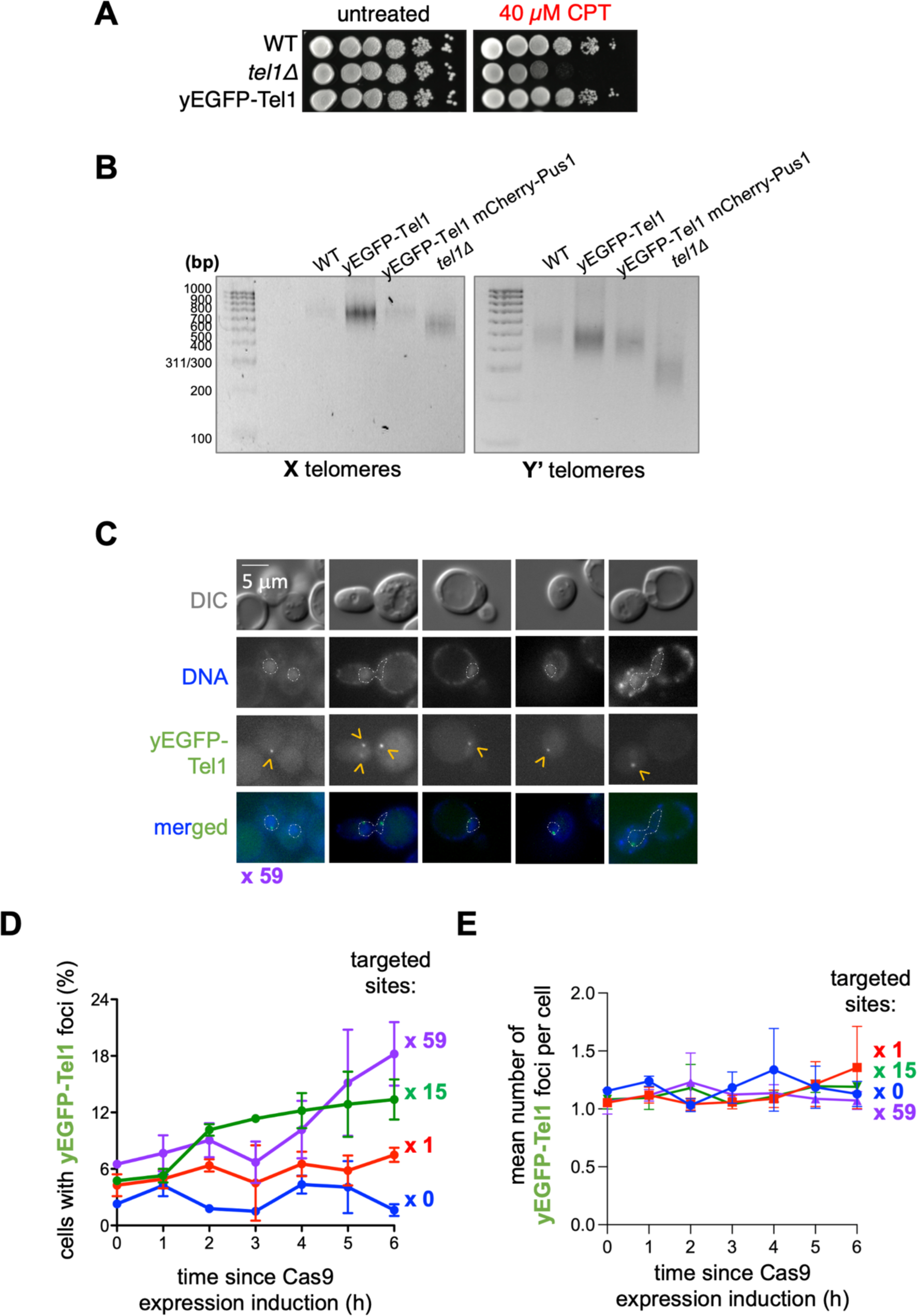
Characterization by microscopy of Tel1 foci formation in response to DSBs. **A.** 10-fold serial dilutions of *S. cerevisiae* cells of the indicated genotypes spotted onto YPED rich medium plates supplemented either with DMSO (untreated) or with 40 µM CPT, incubated 2 days at 30°C and imaged. **B.** Telomeres (X and Y’) length was measured by PCR-mediated amplification (see M&M) from genomic DNA extracted from the indicated strains. *tel1Δ* cells were included as a control for their telomere shortening phenotype. yEGFP-Tel1 cells, whether additionally bearing mCherry-Pus1 or not, display wild-type-length telomeres. **C.** Representative images of yEGFP-Tel1 cells subjected to Cas9-driven DSBs with the x 59 gRNA, as described in (D). The contour of the nucleus has been superimposed using the signals provided by DNA staining with DAPI. yEGFP-Tel1 foci are indicated by yellow arrowheads. **D.** yEGFP-Tel1 cells transformed with the vector expressing an inducible Cas9 and the plasmid expressing the gRNA driving the desired number of DSBs grown to the exponential phase in selective minimal medium using glycerol as the carbon source. Cell samples collected at time 0 before galactose was added to induce Cas9 expression and at the indicated times after induction were inspected by microscopy in search of Tel1 foci. The graph shows the percentage of cells displaying, at least, one Tel1 focus. Each point is the mean of three independent experiments, and the error bars represent the SEM out of those three experiments. At least 150 cells were considered per time, condition and experiment. **E.** The mean number of Tel1 foci per cell (as established by counting the total number of foci divided by the total number of cells) was calculated out of at least 150 cells for each time point and condition. This experiment was done three times, and the plotted value is the mean out of those three experiments. The error bars correspond to the SEM associated to them

Inducing DSBs with the CRISPR-Cas9 dose-dependent system led to Tel1 forming nuclear foci (Fig 6C) whose round morphology and size did not differ from those reported after DSBs induction with ionizing radiations (31), or other DNA damage-related foci, for example of Rad52/Rfa1 (Fig 4A). The basal level of cells presenting Tel1 foci in the population was low, at around 4% (Fig 6D), suggesting that the basal localization of Tel1 at telomeres does not lead to foci formation. The progressive accumulation of DSBs, dependent on time and on gRNA type, matched a parallel increase in the percentage of cells in the population displaying Tel1 foci (Fig 6D, x 0 < x 1 < x 15 < x 59). The maximum obtained value of 20 % (x 59 targeted sites at 6 h) was in striking concordance with the percentage of cells displaying Rad52 foci in the same condition. However, the increase of Tel1 foci happened earlier than the one of Rad52 foci (compare with Fig 4C), hinting at Tel1 role in damage sensing. Finally, positive cells rarely displayed more than one Tel1 focus, even after 6 hours of induction of the x 59 system (Fig 6C, one rare example of three foci per nucleus has been included; quantification in Fig 6E). We conclude that N-terminal tagging of Tel1 with yEGFP represents a performant tool allowing to monitor early events of DSB repair in agreement with its described role in the first steps of DSB signaling.

### Tel1 can form multiple foci at the nuclear envelope upon induction of DSBs

We noted that we always observed Tel1 foci in the periphery of the nucleus (Fig 6C). To know whether this characteristic was specific to DSBs induced by Cas9 at Ty elements, we compared Tel1 foci formation upon Cas9-induced DSBs to chemical induction of DSBs by the clastogen zeocin (41). We grew cells in the same conditions as for the Cas9-DSB-inducing system experiments. We acquired images of these untreated cells and then at intervals till 100 min of exposure to 100 μg/mL zeocin. We could observe that Tel1 foci formed at the same frequency in the population as when DSBs were induced by Cas9, although much faster (Fig 7A). A maximum value of 24 % was reached at 40 min that was invariably maintained till 100 min. Monitoring the percentage of cells bearing Rad52-YFP foci demonstrated an only-slightly delayed kinetics (Fig 7B). Unexpectedly, we observed that zeocin induced a much higher number of Tel1 foci per cell than Cas9. In fact, we observed that the frequency of cells with more than one (up to 8) Tel1 focus per nucleus increased as time passed by under zeocin treatment (Fig 7C). This could be due to zeocin triggering many more DSBs than Cas9 per genome at any given moment. Alternatively, though not exclusively, we reasoned that Ty elements are often inserted near tRNA genes, and that tRNA genes have been described to be clustered in the nuclear space by condensin (42). Single Tel1 foci observed with the Cas9 system could thus relate to DSB clustering by condensin. To test this possibility, we used the thermo-sensitive condensin mutant *smc2-8*, which has been shown to disperse the clusters of tRNA genes (42). Our results show that the proportion of cells containing Tel1 foci increased in the *smc2-8* mutant bearing the x 59 system upon Cas9 induction when cultured at the restrictive temperature of 37°C compared to the permissive temperature of 24°C (Fig 7D). Specific comparison at Cas9-expressing time-points of the percentage of cells displaying 2 Tel1 foci or more demonstrated a significant increase at such restrictive temperature (Fig 7E). Together, these data indicate that the difference in Tel1 foci number following treatment with zeocin compared to CRISPR-guided DSBs is due, at least in part, to the particular spatial organization of Ty elements in the nucleus.

**Figure 7.**
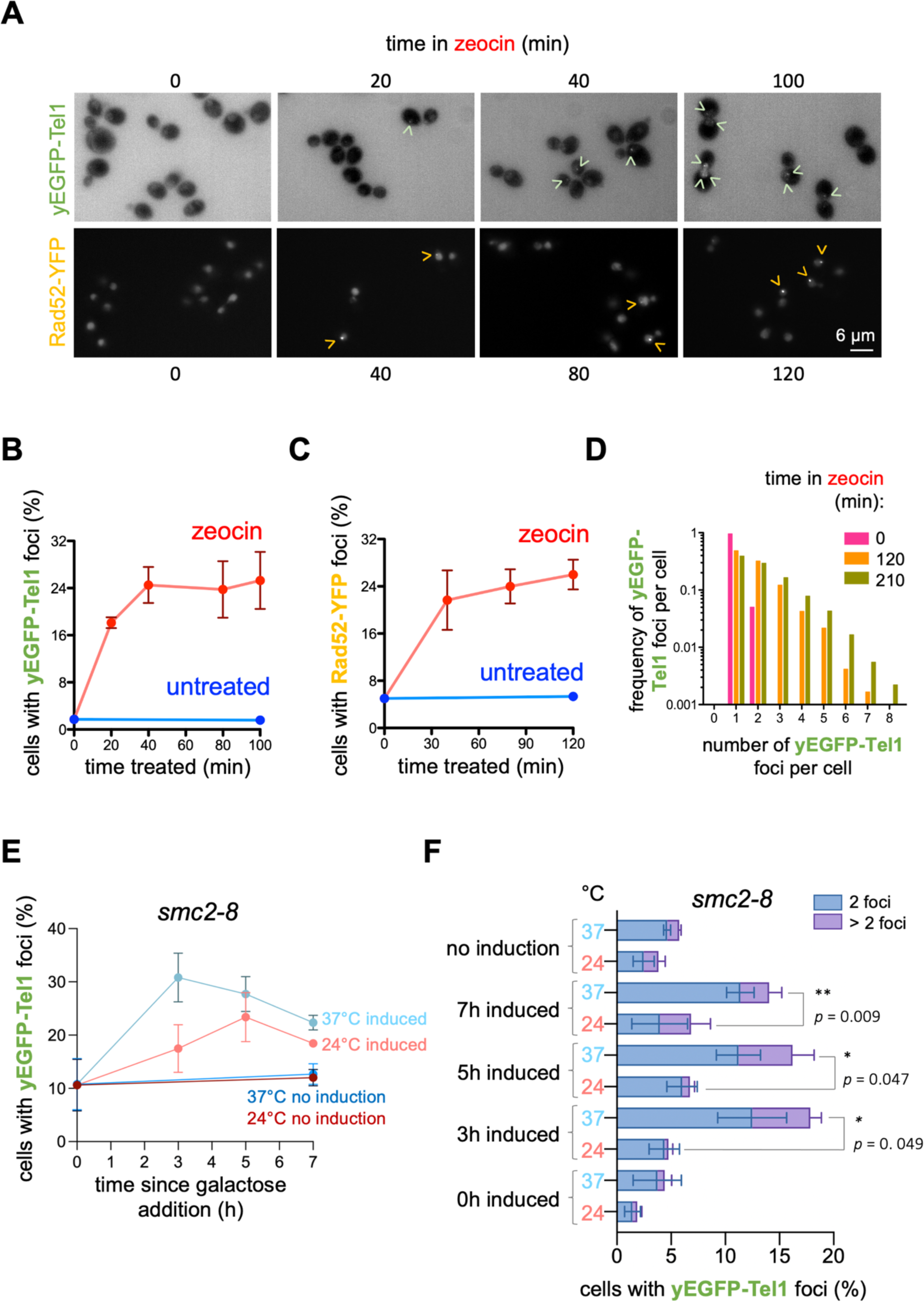
Tel1 can appear in the form of more than one focus per nucleus. **A.** An otherwise wild-type strain tagged with yEGFP at the N-terminus of Tel1 and with mCherry at the N-terminus of Pus1 (to visualize the nucleoplasm) was transformed with an empty vector allowing its growth in minimal selective medium. Cells were exposed to 100 μg/mL zeocin and samples retrieved for analysis by fluorescence microscopy at the indicated times. The graph shows the percentage of cells in the population displaying at least one focus of yEGFP-Tel1 at different times. Each point is the mean of three independent experiments, and the error bars represent the SEM out of those three experiments. At least 150 cells were considered per time, condition and experiment. **B.** Details as in (B) but to score the formation of Rad52-YFP foci. **C.** The foci count data obtained from the cells harbouring at least one focus presented in the three experiments described in **(B)** were exploited to build a frequency distribution. In brief, the number of Tel1 foci per nucleus was counted and a frequency histogram was drawn. Since the three independent experiments provided similar profiles, all the values were merged to build a more robust distribution. The graph illustrates the frequency at which a nucleus displays the indicated number of Tel1 foci for cells not being exposed to zeocin, or exposed for 120 or for 210 minutes (pink, orange and green bars, respectively). **D.** *smc2-8* yEGFP-Tel1 cells bearing the x59 gRNA and inducible Cas9 vectors were grown to the exponential phase in selective medium plus glycerol, to prevent Cas9 expression. The culture was then split, and half set at 24°C, half at 37°C, for 2h. Then, each culture was again split into two, of which one half remained unprocessed (no cut), the other half to which 2% galactose final was added to induce Cas9 expression and genome cleavage. Images to monitor yEGFP-Tel1 foci formation were acquired at the indicated time-points, and the percentage of such foci calculated for three independent experiments (mean and SEM are shown). **E.** The percentage of cells from (E) displaying two or more yEGFP-Tel1 foci is presented as a bar plot for each given condition. The total percentage of the addition of these categories was compared at each meaningful time-point between the permissive (24°C) and the restrictive (37°C) temperatures using a paired *t*-test analysis, as indicated.

Interestingly, the higher number of Tel1 foci induced by zeocin reinforced the earlier observation that they always form near the nuclear periphery (Fig 8A). To investigate further the tight proximity to the nuclear envelope, we simultaneously monitored Tel1 foci and the nucleoporin Nup57, marked in its C-terminus with a tDIMER-RFP moiety. Tel1 foci were recurrently found close to Nup57-tDIMER signals (Fig 8B), with a maximum distance between the maximal intensities of both adjacent signals close to the resolution limit, 0.2 µm (Fig 8C).

**Fig 8.**
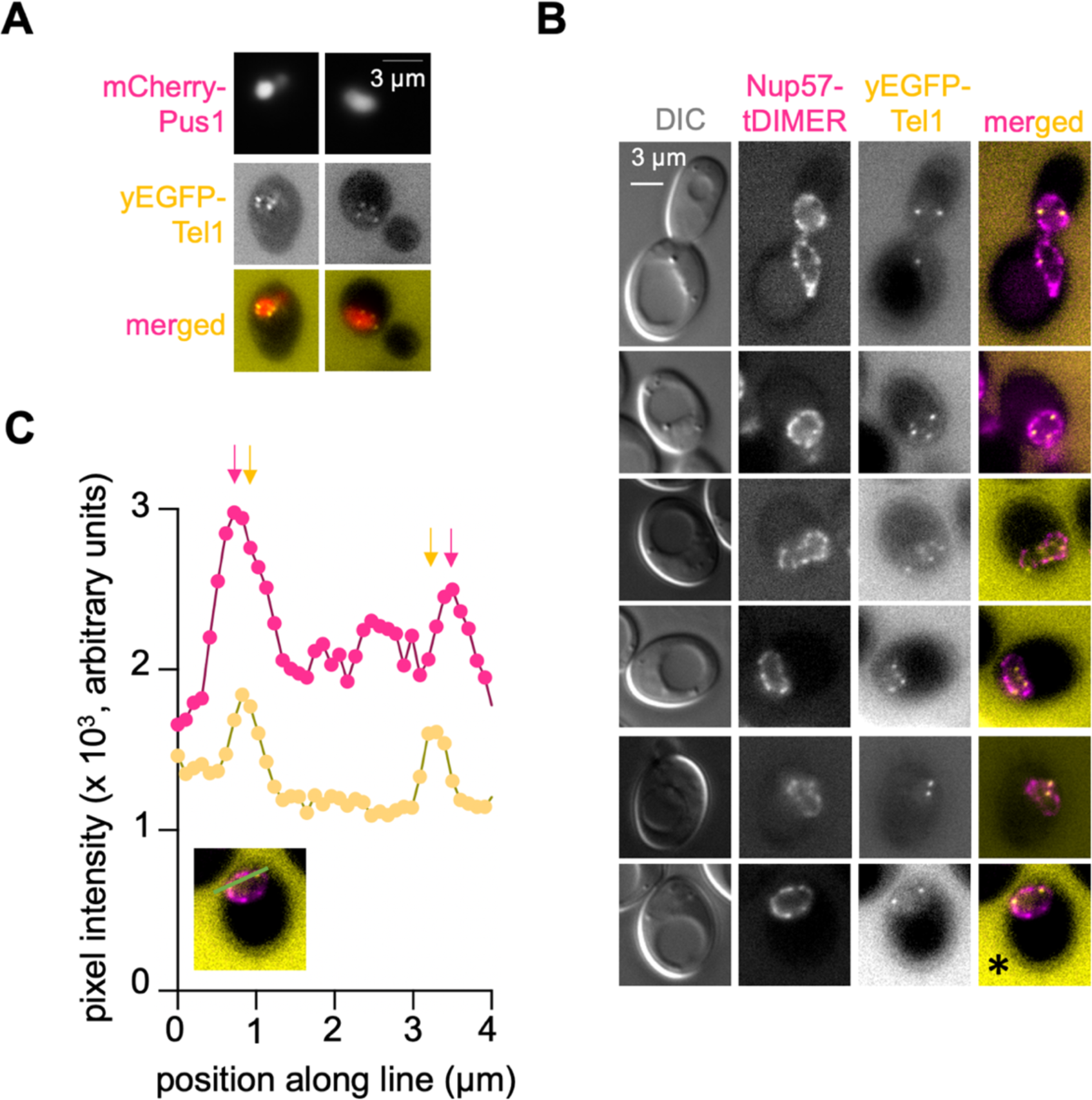
Characterization of the proximity of Tel1 foci to the nuclear envelope. **A.** An otherwise wild-type strain was tagged with yEGFP at the N-terminus of Tel1 and with mCherry at the N-terminus of the nucleosoluble protein Pus1 in order to define the nucleoplasm. The subcellular localization of Tel1 was assessed by fluorescence microscopy in response to 100 μg/mL zeocin. Images of both channels as well as their merging are shown. **B.** An otherwise wild-type strain was tagged with yEGFP at the N-terminus of Tel1, and with tDIMER-RFP at the C-terminus of the nucleoporin Nup57 with the goal of defining the nuclear periphery. The relative position of Tel1 foci with respect to nucleoporin signals was assessed by fluorescence microscopy after exposing the cells for 2 h to 100 μg/mL zeocin. Images of the Differential Interference Contrast (DIC), RFP and GFP channels, as well as their merge are shown. The asterisk marks the cell used to create the graph shown in **(C)**. **C.** A straight line (indicated in green color) was drawn from left to right onto the chosen image and the pixel intensity along it plotted for both the Nup57-tDIMER and yEGFP-Tel1 images. The vertical arrows indicate the points of maximal intensity, thus highlighting the proximity of Tel1 signals to the nuclear periphery.

Altogether, we report here that yEGFP-Tel1 represents a tool to monitor the very early steps of DSB signaling, irrespective of whether they are enzymatically or chemically induced. Furthermore, the subnuclear distribution of Tel1 foci near the nuclear envelope, which is different from the distribution of Rad52-YFP or Rfa1-CFP foci, suggests the existence of yet unknown aspects of initial DSB processing in the nuclear space.

## Discussion

In this work, we have expanded the toolbox in the field of DSB sensing and repair. First, we have taken advantage from the repetitiveness of the transposon elements in the genome of *S. cerevisiae* to design gRNAs capable of driving Cas9 action at specific sites in the genome. Upon Cas9 expression induction, we can compare otherwise identical genomes being broken at an increasing number of locations by the same enzyme. We have validated genetically, physically and functionally the performance of this system. Second, we used this tool to characterize the behavior of the apical kinase of the DNA damage response Tel1 in space and time. We found that Tel1 molecules congregate in the shape of foci clustered by condensin in response to Cas9-induced DSBs in transposon elements. We also show that zeocin-induced DSBs can lead to the formation of up to 8 Tel1 foci per cell, which distribute in tight contact with the nuclear periphery.

Previous works have used the repetitiveness of the Ty elements in *S. cerevisiae* genome to insert 2, 7 or 11 restriction sites that can be cut upon controlled induction of the HO endonuclease (15,16). While the design behind our system is reminiscent of this one, we managed to devise a wider range of induced DSBs, thus permitting further studies on the dose-dependency. Indeed, the Ty-HO system was used to assess, by Southern blot, the role of Mre11 on resection as well as, by monitoring the phosphorylation of the downstream effector Rad53, the role of Tel1 in DNA damage signaling (15,16). Given the sensitivity of Southern and western blotting, a maximum of 7 cuts was enough to assess functional differences. Yet, our system provides an enlarged palette of induced DSBs suitable for less sensitive studies or to extend the study of DSB repair genome-wide. 60% of the human genome being composed of transposable elements, targeting these repetitive elements with the CRISPR-Cas9 technology, as we performed in the yeast genome, would provide a useful tool to produce enzymatic dosedependent DSBs in human cells.

More than 13 proteins working in the cascade of DSB sensing and repair have been fluorescently labelled and their *in vivo* foci formation ability scored by microscopy (31,35,43). With the exception of post-Spo11 cutting during meiosis, with up to 15 Rad52 foci measured per cell (44), these proteins mostly gather in a single focus irrespective of the number of DSBs. For example, even at doses as high as 160 krad, which induces up to 80 DSBs per cell, haploid *S. cerevisiae* cells as the ones used in this study eventually form a maximum of 2 Rad52 foci (44). Of all this set, the only protein openly reported to simultaneously form multiple nuclear foci is Rfa1 (45). The difference in whether the factor gathers under a single focus or as multiple foci may come from the nature of the lesion to be repaired, the DNA-binding properties of the factor under consideration, or other physical properties ruling its ability to phase-separate (45,46). A striking finding of our study is the evidence that Tel1 can form multiple (up to 8) foci per cell upon zeocin addition. Given that Tel1 exerts an early and main role in damage sensing, prior to any engagement of DNA repair activities, the notion arises of whether lesion recognition and lesion repair concur under different regimes of protein nucleation. For example, lesion recognition could occur through a gathering process restricted locally, thus giving rise to more detectable events. Later on, from the moment resection by nucleases would take place, the subsequent steps would be ruled by different nucleation abilities (47).

Analysis of numerous nuclei imaged from distinct angles led us to propose that Tel1 foci form at the periphery of the nucleus. A deeper analysis of Tel1 foci proximity with the nuclear periphery by visualizing a fluorescently tagged nucleoporin further supported this notion. The fact that Tel1 foci form in close contact to the nuclear periphery raises the possibility that a nucleation factor presumably exists close to, or even at, the nuclear membrane that serves to scaffold them. It also suggests that the nuclear periphery could be implicated in DSB sensing. Tel1 and the other DNA damage response apical kinase, Mec1, are Phosphatidyl Inositol 3-kinase-like kinases. Although these kinases are thought not to bear the ability to phosphorylate phosphatidylinositol (PI) moieties any longer (48), they have kept their ability to interact with such molecules. Indeed, the Mec1 human homolog, ATR, was reported to be assisted by phosphoinositides in order to correctly nucleate in the shape of foci upon DNA damage (45). Furthermore, ATR demonstrates ability to sense lipids at membranes (49), and to act at the nuclear membrane to phosphorylate its targets in response to mechanical cues (50). Moreover, Tel1 has been recently reported to bind various lipid moieties inserted within membranes (51), making it very tempting to suggest that Tel1 is being guided at lipid hotspots at the inner nuclear membrane either to exert its DNA breaks sensing activity, or to regulate downstream steps of DSB repair. In a further attempt to venture into this direction, we note a striking similarity of the Tel1 foci arrangement and the discrete spots described along the nuclear-vacuole junction marked by the fatty acids metabolism-related enzyme Mdm1 (52). It will be worth exploring in a near future whether this Tel1 nucleation also relates to the metabolism of lipids.

## Acknowledgements

We thank Philippe Pasero for strains, Olivier Gadal for the vector harboring *NUP57*-tDIMER, and Symeon Siniossoglou for the vector permitting mCherry-Pus1 tagging. We thank Pascale Lesage and Emmanuelle Fabre for insightful comments to improve the manuscript. We acknowledge the imaging facility MRI, a member of the national infrastructure France-BioImaging, supported by the French National Research Agency (ANR-10-INBS-04, Investissements d’avenir). We are thankful to the joint IGMM-CRBM ‘yeast media and technologies service’ for providing us with ready-to-use media. M.M.-C. thanks the ATIP-Avenir program, La Ligue contre le Cancer, and l’Institut National du Cancer (PLBIO19-098 INCA_13832), France, for funding the research in her laboratory. B.P. thanks the Fondation ARC pour la Recherche sur le Cancer (ARCPJA22020060002119) for supporting his work.

## Author Contributions

**Conceptualization:** B.P., M.M.-C.; **Data curation:** J.C., S.K., O.S., B.P., M.M.-C.; **Formal Analysis:** J.C., S.K., O.S., B.P., M.M.-C; **Funding acquisition:** B.P., M.M.-C.; **Investigation:** J.C., S.K., O.S., B.P., M.M.-C.; **Methodology:** S.K., B.P., M.M.-C.; **Project administration:** B.P., M.M.-C.; **Resources:** B.P.; M.M.-C.; **Supervision:** M.M.-C.; **Visualization:** B.P., M.M.-C; **Writing – original draft:** B.P., M.M.-C.; **Writing – review & editing:** J.C., O.S., S.K., B.P., M.M.-C.

## Abbreviations

**CFP**, Cyan Fluorescent Protein; **CPT**, camptothecin; **CRISPR**, Clustered Regularly Interspaced Short Palindromic Repeats; **DIC**, differential interference contrast; **DSBs**, Double Strand Breaks; **gRNA**, guide RiboNucleic Acid; **G_1_**, Gap 1 phase; **G_2_**, Gap 2 phase; **HR**, Homologous Recombination; **NHEJ**, Non-Homologous End Joining; **PAM**, protospacer adjacent motif; **PFGE**, pulsed field gel electrophoresis; **RFP**, red fluorescent protein; **S**, DNA synthesis phase; **WT**, wild type; **yEGFP**, yeast Enhanced Green Fluorescent Protein; **YFP**, Yellow Fluorescent Protein; **YNB**, yeast nitrogen base; **YEPD**, yeast extract peptone dextrose.

## Supporting Information

**S1 Fig.**
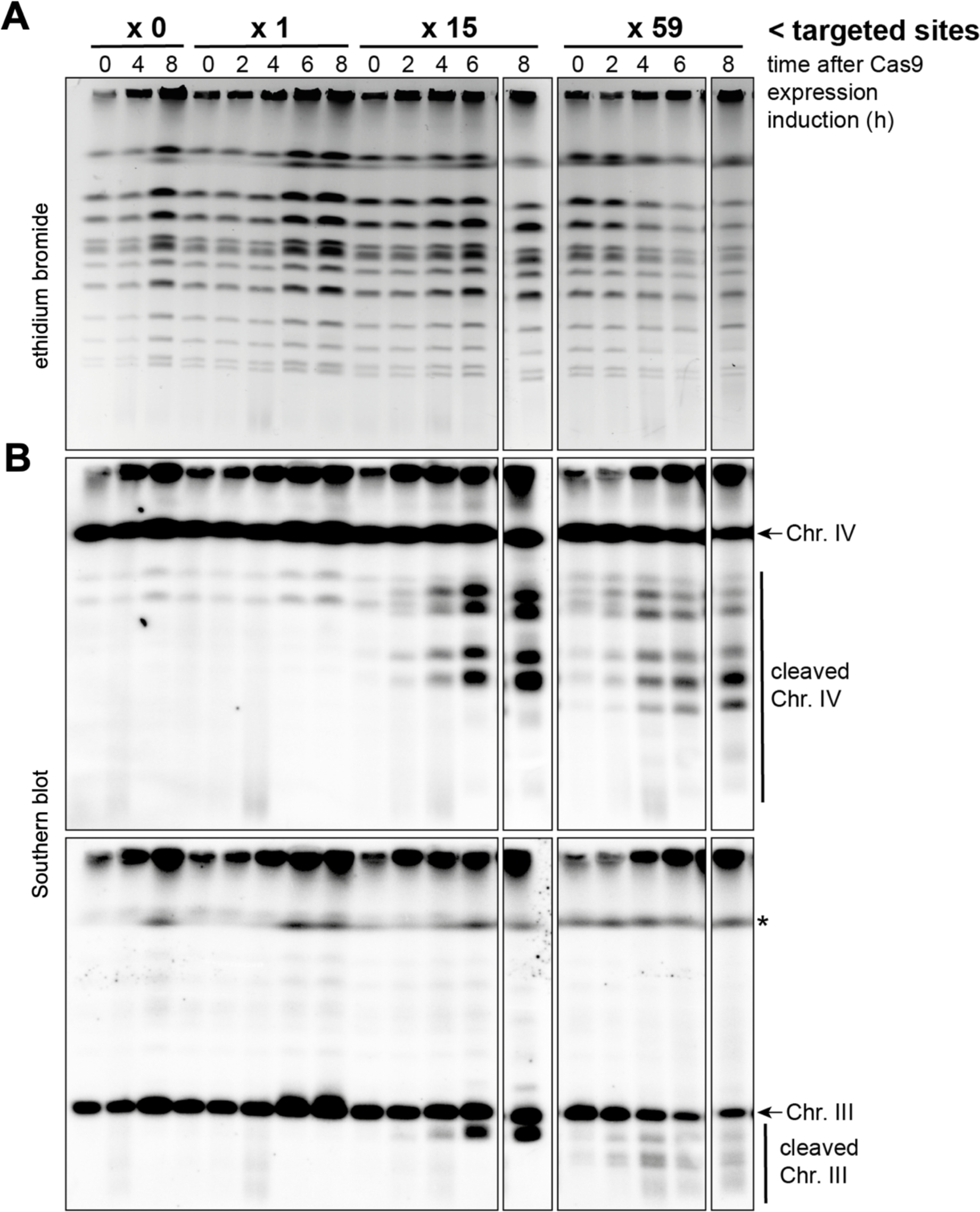
PFGE used for restriction analyses of chromosome cleavage by Cas9. **A.** PFGE was prepared and run as in **Fig 2B**, and stained with ethidium bromide. Please note that the time point 8 h for the x 15- and x 59-cuts were inadvertently exchanged during gel loading. As such, the broken boxes indicate that the two lanes have now been replaced where they belong. **B.** Southern blot hybridizations against chromosome (Chr.) IV (top) and III (bottom) against the DNA run in **(A)**. The asterisk denotes a residual band after incomplete stripping of chromosome IV hybridization.

**S2 Fig.**
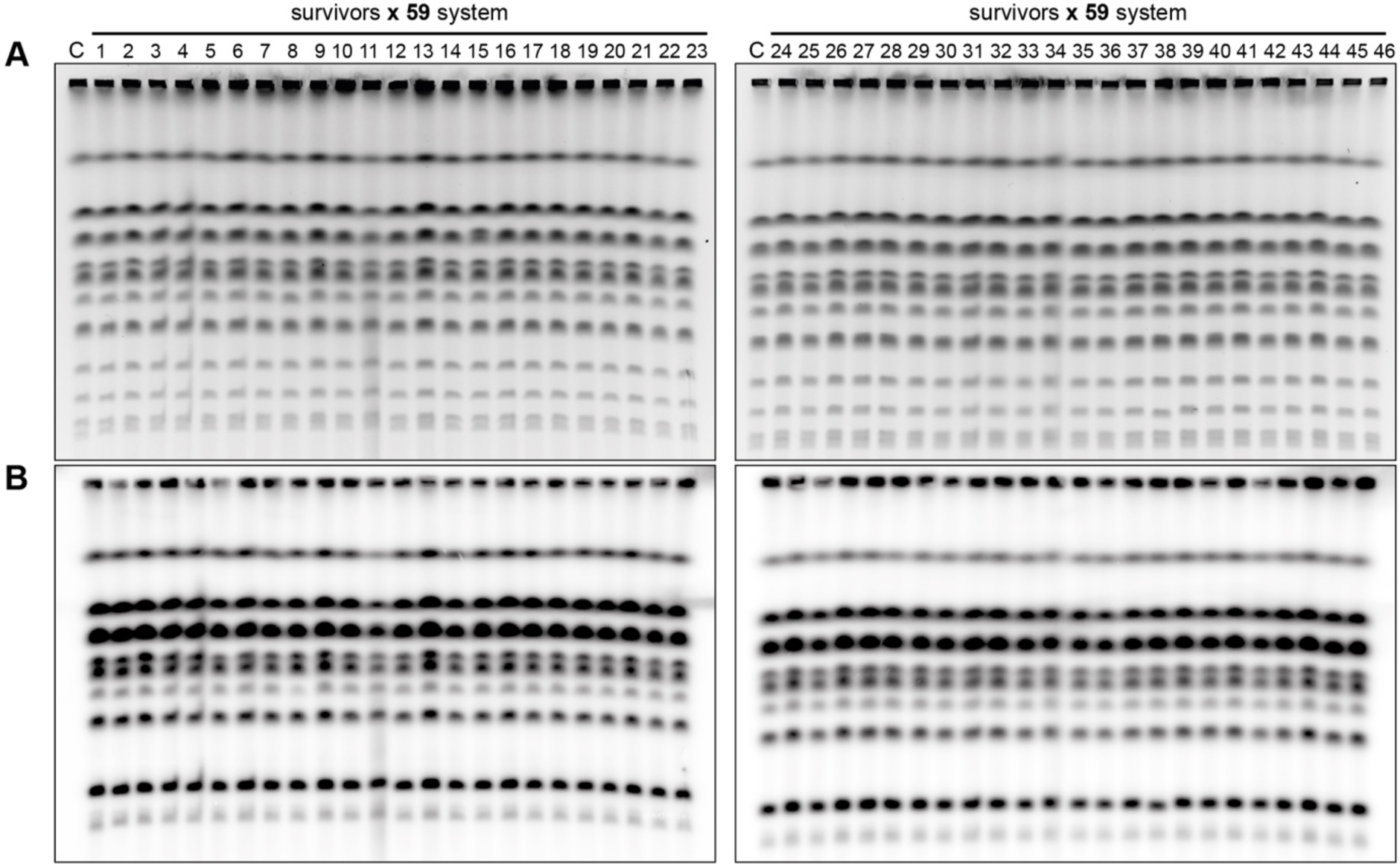
Analysis of chromosome integrity after DSB repair. Cells bearing the Cas9 and the x59-cuts gRNA plasmids were first exposed to galactose as to induce Cas9 expression for 8 h, then seeded onto rich medium plates containing glucose, in order to repress Cas9 expression and allow survival following DSB repair. 23 colonies from two independent experiments were used to prepare DNA in plugs and assess survivors’ karyotype by Pulsed Field Gel Electrophoresis (PFGE). **A**, ethidium bromide gels. **B**, Southern blot after these electrophoreses using a probe specific for Ty1 and Ty2 elements. C: karyotype from cells taken before Cas9 expression induction.

